# Prescribed Burns Drive Lasting Changes in Soil Nitrogen Cycling and Microbial Function

**DOI:** 10.64898/2026.01.21.700861

**Authors:** Alaina O. Benot, Gray Waldschmidt, Isaac J. Okyere, Eva O.L. Legge, Andrew L. Vander Yacht, Samuel C. Gilvarg, Chanistha Tiyapun, Jennifer L. Goff

## Abstract

Fire can be a major pulse disturbance to soil microbial communities. Yet regular burning is a natural and essential process that maintains biodiversity and the unique attributes of rare and imperiled fire-dependent ecosystems. Most studies of fire effects on soil microbial communities typically focus on short-term (<1 year) responses following a single fire event. Here we examined the longer-term effects of repeated prescribed fire at the Albany Pine Bush—a fire-dependent, inland pitch pine (*Pinus rigida*) barren ecosystem of the northeastern US. We observed that this long-term fire management (*i*.*e*., a fire interval of approximately every 5 to 13 years over the past 30 years) has led to substantial depletion of soil nitrogen, specifically nitrate. However, we found no lasting shifts in the higher-level taxonomic composition of soil prokaryotic communities. Instead, metagenomic analysis revealed significant changes in the nitrogen-cycling functional potential, specifically, decreased dissimilatory nitrate reduction and denitrification potential in repeatedly burned soils. Collectively, these data suggest fire-induced geochemical changes persist under repeated burning, potentially driving substantial shifts in soil microbial functional diversity. Our study reveals strain-level changes that would be missed when examining only higher-level taxonomic patterns. Where fire is repeatedly applied, fire-induced shifts in soil microbial communities can persist well beyond a few weeks after burning—challenging prevailing views of short-lived belowground effects of prescribed burns.

## Brief Communication

Ecological disturbances, such as fire, alter microbial communities and their activities within Earth’s biogeochemical cycles [1–3]. Yet, research on fire-microbe feedbacks has overwhelmingly focused on short-term responses to single, high-intensity wildfires, typically within the first year post-burn [4,5]. Despite the increasing use of prescribed fire (*i*.*e*., an intentionally burned area) to mitigate wildfires and foster biodiversity, the long-term impact of repeated prescribed fires on soil microbial communities remains poorly understood. Available evidence suggests that taxonomic shifts following a single prescribed fire are modest and transient, often dissipating within weeks to months [3,6–8]. Whether repeated burning produces persistent structural and functional changes remains unclear. Given prescribed fire’s impact on aboveground biodiversity, we hypothesized that multi-decadal fire management imprints a lasting geochemical signature in soil that fundamentally reshapes microbiome functional potential, with far-reaching implications for nutrient cycling and trace gas fluxes [9].

Although the use of fire by Indigenous communities in the region dates back millennia [10], the Albany Pine Bush Preserve (APBP, Albany, N.Y., U.S.A. [42.71 °N, 73.86 °W]) has a contemporary prescribed burn history dating back approximately 30 years [11,12]. This fire-dependent landscape has a pitch pine (*Pinus ridiga*) dominated canopy with a scrub oak (*Quercus ilicifolia*), dwarf oak (*Q. prinoides*), and herbaceous plant understory [12]. Fire maintains nutrient-poor soils, preventing succession into closed-canopy hardwood forests [12,13]. Thus, the APBP serves as a powerful model system for examining long-term impacts of prescribed fire on soil microbiomes.

We sampled soils from nine stands at the APBP, spanning a gradient of prescribed fire frequency over the past 30 years: “frequent” stands experienced ≥4 fires (average fire return interval = 5.7 years) while “infrequent” stands experienced >1 but ≤3 fires (average fire return interval = 13.3 years), reflective of the high and low ends, respectively, of historic fire return intervals [14]. “Control” plots had no record of fire over this period **(Fig. 1A, Tables S1-S2)**. Within each stand, soils were collected at four sampling sites from two depths (0-15 cm and 15-30 cm), homogenized by depth and stand, and stored at −80°C before analysis (n = 18 samples) **(Table S3)**. Metagenomic sequencing and assembly methods are described in Benot et al. (2025) [15]. Taxonomic profiles were generated using Kaiju, and nitrogen-cycling genes were quantified using Fama, both implemented on the DOE KnowledgeBase Platform [16]. Soil geochemical analyses **(Table S4)** included: pH, % organic matter, % total nitrogen (TN), % total carbon (TC), exchangeable acidity, effective cation exchange capacity, ammonium chloride-extractable-calcium, potassium, magnesium, phosphorus, aluminum, iron, manganese, sodium, and zinc, nitrate (NO_3_^−^-N), and ammonium (NH_4_-N). For additional details, see **SUPPORTING METHODS**.

**Figure 1.**
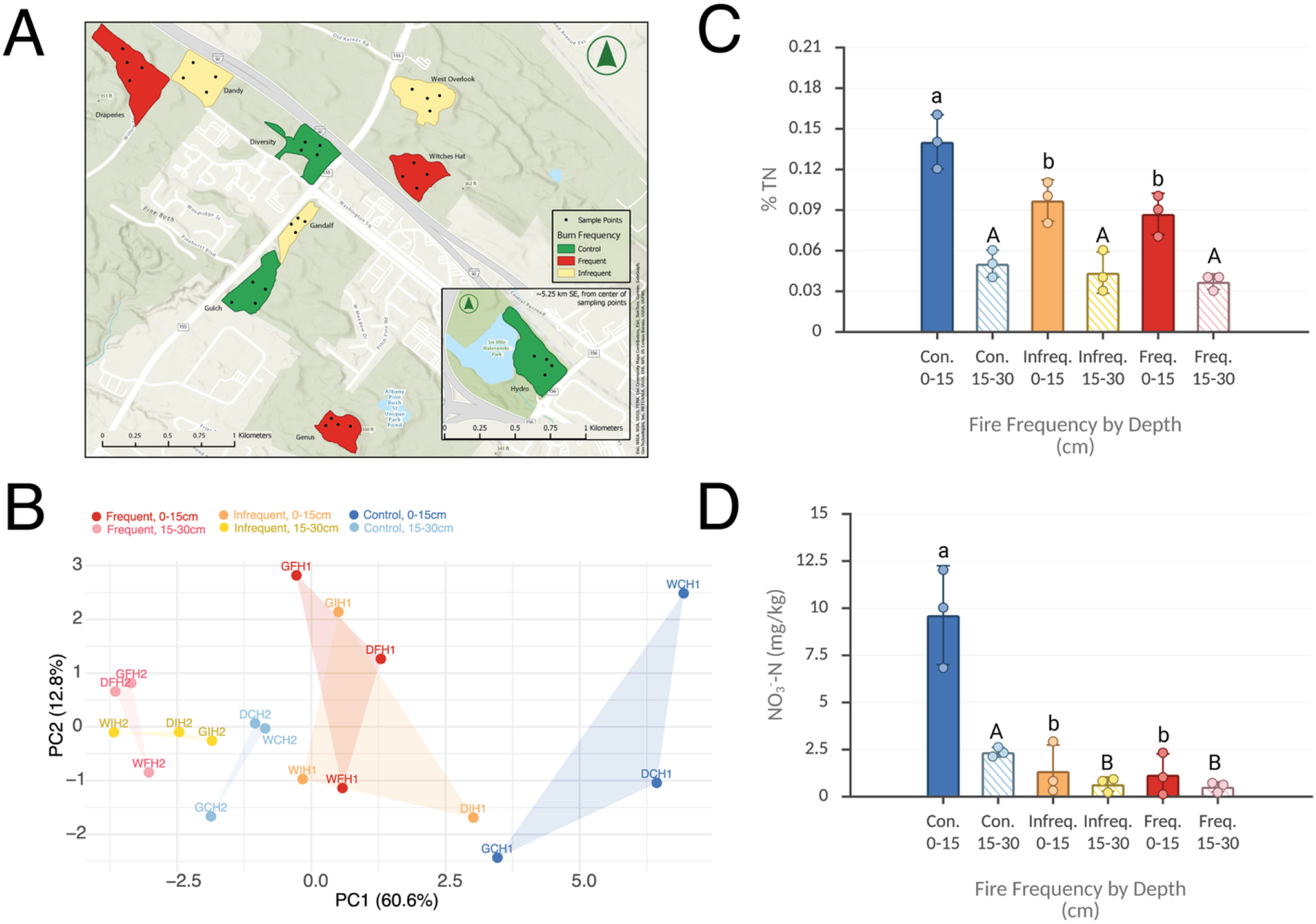
**A**: Location of randomly distributed sub-sampling sites within selected APBP stands. **B**: PCA plot of homogenized sample site geochemistry. **C** and **D**: Bar graph of average NO_3_^−^-N (mg/kg) and average percentage total nitrogen, respectively. One-way ANOVAs were conducted independently for each soil depth, followed by a post hoc Tukey’s HSD test. Lowercase (a, b) and uppercase (A, B) letters indicate significantly different (*p* < *0*.*05*) treatment means at 0-15 cm and 15-30 cm depths, respectively. Bar graphs were created in https://BioRender.com.

Fire affects the soil surface [13]. Yet geochemical changes observed at both soil depths suggested both acute and sustained impacts of fire on soil chemistry **(Table S5**). Principal component analysis (PCA) of the geochemical data **(Table S4)** revealed separation of sites by both fire management history and depth **(Fig. 1B)** that involved significant shifts in soil nitrogen (N) chemistry. At 0-15 cm, %TN decreased ~1.5-fold in both fire treatments compared to controls (one-way ANOVA with Tukey HSD, *p* < *0*.*05* for both treatments) **(Fig. 1C)**. No differences in NH_4_-N were observed between treatments (one-way ANOVA, *p* > *0*.*05*). However, bioavailable NO_3_^−^-N declined substantially with burning at both depths, with 9-fold (frequent) and 5-fold (infrequent) reductions in the surface soils and similar declines at the 15-30 cm depth (5-fold and 4-fold, respectively) (one-way ANOVA with Tukey HSD, *p* < *0*.*05*) **(Fig. 1D)**. Possible mechanisms for this NO_3_^−^-N loss includes incorporation into recalcitrant pyrogenic organic matter [4,17] and volatilization [18].

Subsequently, read-based taxonomic mapping was used to evaluate whether fire-driven soil changes altered microbial community composition. Sample depth had a significant effect on community composition from the phylum to genus levels (PERMANOVA, *p* < *0*.*05*), whereas fire treatment and its interaction with depth were not significant influences at these taxonomic levels (PERMANOVA, *p* > *0*.*05*). This aligns with prior studies reporting minimal or no detectable change in community composition weeks to months after a prescribed fire [8,19]. Across all soils, the most prevalent phyla were *Proteobacteria*, followed by *Acidobacteria* and *Actinobacteria* (**Fig. S1, Tables S6-S10)**.

Given significant shifts in soil N in fire-managed sites, the metagenomic reads were mapped to microbial N-cycling genes **(Table S11)**. Unlike the community analysis, variation in N-cycling gene composition was significantly explained by both fire treatment and depth (PERMANOVA, *p = 0*.*004* and *p = 0*.*005*, respectively)—with treatment as the strongest predictor (25.5% of variance explained) **(Fig. 2A)**. These are likely species- or strain-level differences not resolved by coarser taxonomic analyses [20]. Depleted bioavailable NO_3_^−^-N in fire-managed soils may be one driver of these shifts. Indeed, there was a strong positive correlation (Pearson’s R > *0*.*60*) between soil nitrate concentrations and the abundances of dissimilatory nitrate reduction and denitrification genes **(Table S12)**.

**Figure 2.**
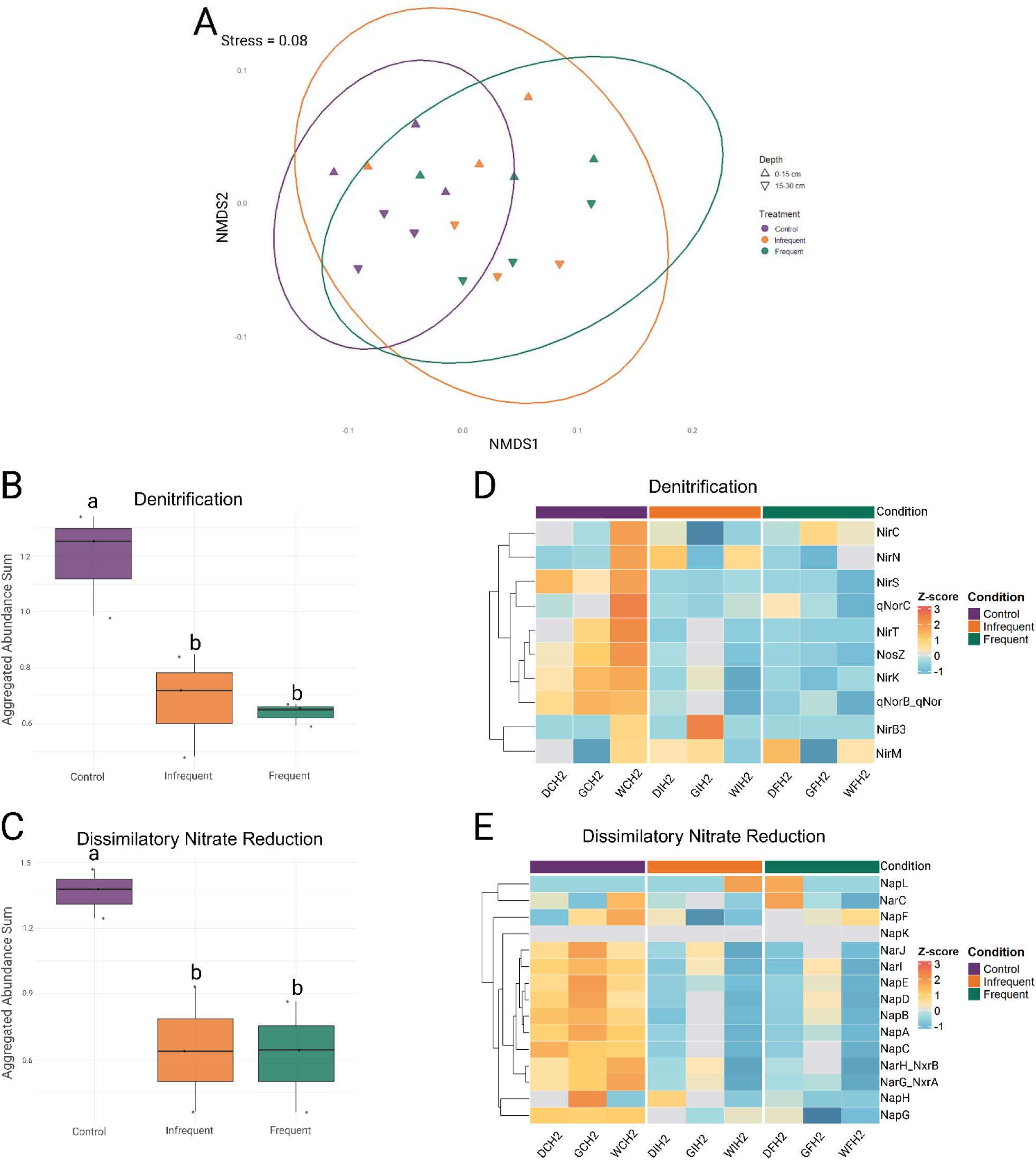
**A:** Nonmetric multidimensional scaling using Bray-Curtis dissimilarity of N-cycling genes in individual samples. **B:** Normalized heatmap of individual genes in the denitrification pathway. **C:** Normalized heatmap of individual genes in the dissimilatory nitrate reduction pathway. For **B** and **C**, one-way ANOVAs were conducted, followed by a post hoc Tukey’s HSD test, and lowercase letters (a, b) indicate significantly different (*p*<*0*.*05*) treatment means. **D:** Gene abundance of dissimilatory nitrate reduction pathway genes at 15-30 cm grouped by treatment. **E:** Gene abundance of denitrification pathway genes at 15-30 cm grouped by treatment. Figure layout was performed in https://BioRender.com.

We also observed significant decreases in the abundances of the denitrification and dissimilatory nitrate reduction pathways **(Fig. 2B-E)** at lower soil depths (15-30 cm) at fire-managed relative to control sites (one-way ANOVA with Tukey HSD, *p* < *0*.*05*). In contrast, the abundances of nitrogen fixation, ammonium oxidation, nitrite assimilation, ammonification, urea degradation, or assimilatory nitrate reduction pathways **(Figs. S3-7)** were unchanged. As these pathways are linked to ammonium flux, the stable ammonium concentrations across treatments may explain the absence of detectable shifts. In contrast, within the denitrification/dissimilatory nitrate reduction pathways, *napA* (encoding periplasmic nitrate reductase), *nirK* (copper-containing nitrite reductase), *qnor* (quinol-dependent nitric oxide reductase), and *nosZ* (nitrous oxide reductase) all decreased in abundance at the lower depth **(Fig. S9)**. At lower soil depths, most mapped reads for *napA, nirK, nosZ*, and *qnor* were associated with *Proteobacteria*, with smaller contributions from *Bacteroidetes* (*napA, nosZ*), *Acidobacteria* (*qnor*), *Actinobacteria* (*nirK, qnor*), *Chloroflexi* (*nirK*), *Nitrospirae* (*nirK*), and *Planctomycetes* (*qnor*) **(Table S13)**. Relative to control soils, fire management reduced relative abundances of *Proteobacteria*-associated *napA* and *nirK*. In contrast, relative abundances of *Proteobacteria*-associated *nosZ* increased, while relative abundances of *nosZ* reads associated with *Bacteroidetes* declined. Together, these shifts indicate that long-term fire management also restructures the taxonomic partitioning of dissimilatory nitrate reduction/denitrification potential.

Despite extensive research on wildfire feedbacks on soil microbiomes, the impacts of long-term prescribed fire regimes on microbial communities remain poorly characterized [3–6]. Prior work suggests that single prescribed burns have limited, short-lived effects on microbial taxonomic composition—relative to more intense and severe wildfire—with detectable changes generally disappearing within weeks to months [6–8,19]. In contrast, our results reveal persistent fire-driven restructuring of microbial functional potential, particularly within N-cycling pathways. Specifically, we observe a sustained reduction of microbial denitrification potential, linking prescribed fire to long-term modulation of soil reactive N gas fluxes—specifically nitrous acid, nitric oxide, and nitrous oxide. Together, these findings demonstrate that repeated prescribed fire restructures soil microbiomes in ways that taxonomic analyses alone cannot detect.

## Acknowledgments

We gratefully thank The Albany Pine Bush Preserve Commission (specifically Neil Gifford, Steve Campbell, Tyler Briggs, and Alex Soldo) for their expertise and guidance regarding the Albany Pine Bush Preserve. We also thank the University of Delaware Ammon-Pinizzotto Biopharmaceutical Innovation Center DNA Sequencing & Genotyping Center for conducting DNA sequencing and the University of Maine for soil geochemical analysis. Also, we’d like to acknowledge Matthew Lathrop for their GIS technical expertise and Aaron Ninokawa for guidance with statistical analyses in R. Financial support for this project came from SUNY ESF startup funds to JLG.

## Data Availability

All geochemical data and site metadata are available in the supplemental information files. All sequencing data used here are associated with BioProject PRJNA1269515. Raw sequencing data are available through NCBI Sequence Read Archive (SRA) via accession numbers SRX28992350–SRX28992355; SRX29211976–SRX29211981; and SRX29333943–SRX29333948. Tables S1, S3-S4, S6-S11, and S13 can be found on Figshare: https://doi.org/10.6084/m9.figshare.31083271.

