## Supplemental Information for "Prescribed Burns Drive Lasting Changes in Soil Nitrogen Cycling and Microbial Function"

**Supporting Methods**

Sampling Methods

Soil Geochemical Measurements

Bioinformatics

Statistical Methods

References

**Supplementary Figures and Tables**

Table S1. Site Metadata (found in external Excel file)

Table S2. Stand Fire History

Table S3. Homogenized Samples (found in external Excel file)

Table S4. Raw Geochemical Data (found in external Excel file)

Table S5. Averaged Soil Geochemistry

Table S6. Kaiju Phyla (found in external Excel file)

Table S7. Kaiju Classes (found in external Excel file)

Table S8. Kaiju Orders (found in external Excel file)

Table S9. Kaiju Families (found in external Excel file)

Table S10. Kaiju Genera (found in external Excel file)

Table S11. FAMA Gene Abundance (found in external Excel file)

Table S12. Correlations Between NO3⁻-N and Gene Abundances

Table S13. FAMA Counts by Phylum (found in external Excel file)

Figure S1. Phylum level prokaryotic community composition

Figure S2. Heatmap and Box-And-Whisker Plot of Genes In Nitrogen Fixation Pathway

Figure S3. Heatmap and Box-And-Whisker Plot of Genes In Ammonium Oxidation Pathway

Figure S4. Heatmap and Box-And-Whisker Plot of Genes In Nitrite Assimilatory Pathway

Figure S5. Heatmap and Box-And-Whisker Plot of Genes In Ammonification Pathway

Figure S6. Heatmap and Box-And-Whisker Plot of Genes In Nitrate Assimilation Pathway

Figure S7. Heatmap and Box-And-Whisker Plot of Genes In Urease Pathway

Figure S8. Heatmap and Box-And-Whisker Plot of Genes In Anerobic Ammonium Oxidation Pathway

***Sampling Methods:*** We established 4 sampling plots within nine 5- to 20-hectare stands at the APBP. Stands were intentionally selected to span variation in the frequency of prescribed fires over the past 30 years (*i.e.*, since 1995): For all selected stands, the most recent fire was at least a full year before our observations. Within each stand, sampling plots were randomly located an average of 44 meters from stand boundaries to minimize edge effects **(Figure 1A)**. At each plot location, we extracted two soil samples – one for each of two soil depths (0-15 cm and 15-30 cm). Samples were then homogenized by soil depth within stands to yield 3 samples per fire treatment and 2 soil depths (total n = 18) and stored at -80 °C until analysis.

***Soil Geochemical Measurements:*** Geochemical analyses were performed at the University of Maine Analytical Laboratory and Maine Soil Testing Service. Soil pH was measured in distilled water. Organic matter was determined by loss on ignition (LOI) at 550°C. Total nitrogen and carbon were measured by combustion analysis at 1350°C. Nitrate and ammonium nitrogen were first extracted in potassium chloride and then measured colorimetrically by Flow Injection Analysis. All other nutrients were extracted in ammonium chloride and measured by inductively coupled plasma optical emission spectroscopy (ICP-OES). Exchangeable acidity was extracted in potassium chloride and measured by titration. Effective cation exchange capacity (ECEC) was calculated by the summation of exchangeable base cations plus exchangeable acidity.

***Bioinformatics:*** Read-based taxonomic mapping was performed using Kaiju (v1.9.0) [1] run on the US Department of Energy’s Knowledge Base (KBase) platform [2]. Read mapping to the nitrogen cycle pathways and individual genes was performed using the Fama pipeline (v1.1) [3], also on KBase. Mapped reads were normalized by library size, target gene effective size, and predicted average genome size. Pathway abundances are inferred from the abundances of their associated functional genes: ammonification (*nrfABCDH*), ammonium oxidation (*amo/pmoABC*), anaerobic ammonium oxidation (*hao, hzo, hzsABC*), denitrification (*nirCKMNST, nirB3, nosZ, cnor, qnor*), assimilatory nitrate reduction (*nasAB*), dissimilatory nitrate reduction (*napABCDEFGHKL, narC, narG/nxrA, narH/nxrB*), nitrite assimilation (*nasIJ, nirABDU*), nitrogen fixation (*anfG/vnfG, nifB, nifD/anfD/vnfD, nifH/anfH/vnfH, nifK/anfK/vnfK*), urease (*ureABC*). The Fama code is freely available through GitHub: <https://github.com/aekazakov/FamaProfiling/tree/d9db15ea217e3be2aab65c356564a6d345b4f410>.

***Statistical Methods:*** Comparisons of soil geochemistry between treatments were performed using a one-way Analysis of Variance (ANOVA) with a post hoc Tukey Honestly Significant Difference (HSD) test conducted using the base R *aov* and *TukeyHSD* functions. A Principal Component Analysis (PCA) was conducted using the base R “stats” *prcomp* function. Comparisons of community composition between treatments were performed using a Permutational Multivariate Analysis of Variance (PERMANOVA) conducted in R using the “vegan” package *adonis2* function (Bray-Curtis distance with permutations = 999) while *metaMDS* was used for Nonmetric Multidimensional Scaling (Bray-Curtis distance matrix) [4]. The “ggplot2” packaged was used for visualizations [5].

| **Table S2. Stand Fire History** | | | | |
| --- | --- | --- | --- | --- |
| **Stand** | **Sample ID Prefix** | **Fire Frequency** | **Total Number of Fires Since 1995** | **Fire Dates*** |
| Draperies | DF | Frequent | 5 | 2005, 2011, 2018, 2023 |
| Witches Hat | WF | Frequent | 4 | 1999 (wildfire), 2005, 2013, 2021 |
| Genus | GF | Frequent | 7 | 1995,1996, 2004, 2004, 2019, 2023, 2023 |
| Dandy | DI | Infrequent | 2 | 2006, 2011 |
| West Overlook | WI | Infrequent | 3 | 1999 (wildfire), 2017, 2022 |
| Gandalf | GI | Infrequent | 2 | 1997, 2022 |
| Diversity | DC | Control | 0 | N/A |
| Hydro | WC | Control | 0 | N/A |
| Gulch | GC | Control | 0 | N/A |

*Unless otherwise marked as a wildfire, all fires were prescribed burns conducted by the APBP

|  | **Table S5. Averaged Soil Geochemistry (AVG ±SD)** | | | | | | | | | | | | | | | | |
| --- | --- | --- | --- | --- | --- | --- | --- | --- | --- | --- | --- | --- | --- | --- | --- | --- | --- |
| **Treatment** | pH | % LOI | % TN | % TC | NO3-N* | NH4-N* | Ca* | K* | Mg* | P* | Al* | Fe* | Mn* | Na* | Zn* | Acidity^^^ | ECEC^^^ |
| **0-15 cm** | | | | | | | | | | | | | | | | | |
| Frequent | 5.3±0.2 | 3.2±0.8 | 0.09±  0.03 | 1.6±  0.5 | 1.1±0.8 | 7.4±5.2 | 234.7±160.1 | 50.3±  28.1 | 20.7±7.4 | 1.4±0.6 | 84.7±  18.0 | 3.8±0.5 | 6.3±3.8 | 4.6±4.7 | 1.1±0.1 | 1.1±0.2 | 2.6±0.7 |
| Infrequent | 5.3±0.2 | 4.1±1.4 | 0.10±  0.02 | 1.7±  0.3 | 1.3±1.4 | 9.8±5.0 | 147.7±90.6 | 40.7±  23.2 | 14.3±5.6 | 2.2±0.8 | 103.7±  40.1 | 4.5±1.3 | 5.6±1.9 | 7.8±0.6 | 1.7±0.4 | 1.3±0.5 | 2.3±0.3 |
| Control | 4.7±0.2 | 4.7±1.2 | 0.14±  0.02 | 2.6±  0.8 | 9.6±2.6 | 8.1±4.3 | 342.0±183.9 | 56.3±  17.8 | 34.7±  15.5 | 3.0±0.6 | 101.7±  23.2 | 7.7±2.6 | 15.0±  7.8 | 12.0±  10.0 | 2.7±1.3 | 1.5±0.1 | 3.7±1.1 |
| **15-30 cm** | | | | | | | | | | | | | | | | | |
| Frequent | 5.4±0.2 | 1.7±0.8 | 0.04±  0.01 | 0.6±  0.2 | 0.5±0.3 | 2.1±0.5 | 50.7±  33.6 | 15.7±  0.6 | 4.3±1.0 | 1.4±1.0 | 51.3±  7.8 | 2.4±0.4 | 1.4±0.9 | 12.5±  17.7 | 0.6±0.9 | 0.7±0.0 | 1.1±0.1 |
| Infrequent | 5.1±0.1 | 1.6±0.3 | 0.04±  0.02 | 0.7±  0.2 | 0.6±0.4 | 3.0±1.7 | 45.3±  23.4 | 26.0±  18.4 | 5.3±1.9 | 1.1±0.2 | 63.3±  12.7 | 3.3±0.7 | 2.4±0.7 | 5.5±2.5 | 0.7±0.7 | 0.8±0.1 | 1.2±0.3 |
| Control | 5.1±0.1 | 2.0±0.6 | 0.05±  0.01 | 0.9±  0.4 | 2.3±0.3 | 2.9±2.1 | 90.3±  30.0 | 36.3±  18.5 | 11.7±  3.2 | 2.1±0.9 | 69.3±  1.5 | 3.4±1.0 | 5.0±3.1 | 11.5±  5.9 | 1.0±0.6 | 0.9±0.0 | 1.6±0.2 |

*mg/kg

^ meq/100 g

Unprocessed data are available in Table S3.

| **Table S12. Correlations Between NO_3_⁻-N and Gene Abundances** | | | |
| --- | --- | --- | --- |
| **Gene Product** | **Pearson's R*** | **Gene Product** | **Pearson’s R*** |
| AmoB_PmoB | 0.162 | NarC | -0.265 |
| HAO | -0.071 | NarH_NxrB | 0.437 |
| **NapA** | **0.659** | **NarI** | **0.611** |
| NarG_NxrA | 0.44 | **NarJ** | **0.746** |
| NasA | 0.217 | NasB | 0.342 |
| NifK_AnfK_VnfK | -0.027 | NasI | -0.088 |
| NirA | 0.209 | NasJ | -0.091 |
| NirB | 0.092 | NifB | -0.103 |
| **NirK** | **0.69** | NifD_AnfD_VnfD | -0.022 |
| NirS | -0.212 | NifH_AnfH_VnfH | 0.091 |
| **NosZ** | **0.656** | NirB3 | 0.273 |
| NrfA | 0.228 | NirC | -0.103 |
| UreC | -0.182 | NirD | 0.185 |
| cNor-C | 0.331 | NirM | -0.372 |
| **cNorB_qNor** | **0.732** | NirN | -0.001 |
| AmoA_PmoA | -0.012 | NirT | 0.149 |
| AmoC_PmoC | 0.188 | NirU | 0.486 |
| AnfG_VnfG | 0.314 | NrfC | -0.158 |
| HzsB | -0.398 | NrfD | -0.107 |
| **NapB** | **0.653** | NrfH | 0.144 |
| **NapC** | **0.645** | UreA | -0.214 |
| **NapD** | **0.647** | UreB | -0.125 |
| NapE | 0.546 | NarC | -0.265 |
| NapF | 0.078 | NarH_NxrB | 0.437 |
| NapG | -0.308 | **NarI** | **0.611** |
| NapH | 0.422 | **NarJ** | **0.746** |
| NapL | -0.226 | NasB | 0.342 |

*Highly positively correlated results (R > 0.60) are bolded

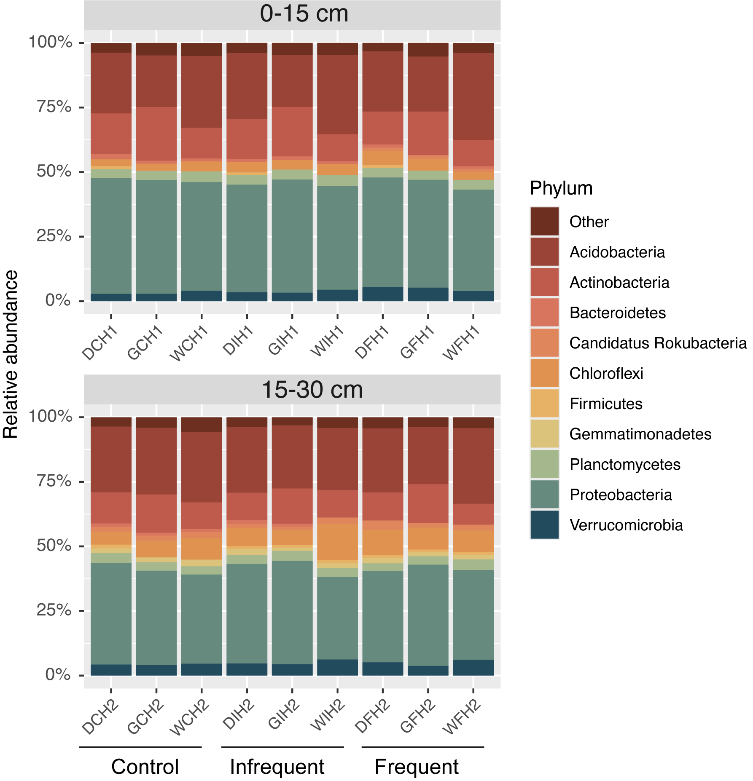

**Figure S1.** Prokaryotic community composition at the phylum level determined through read-based mapping using Kaiju. Compositions for each site are grouped by treatment and shown at both (0-15 and 15-30 cm) soil depths.

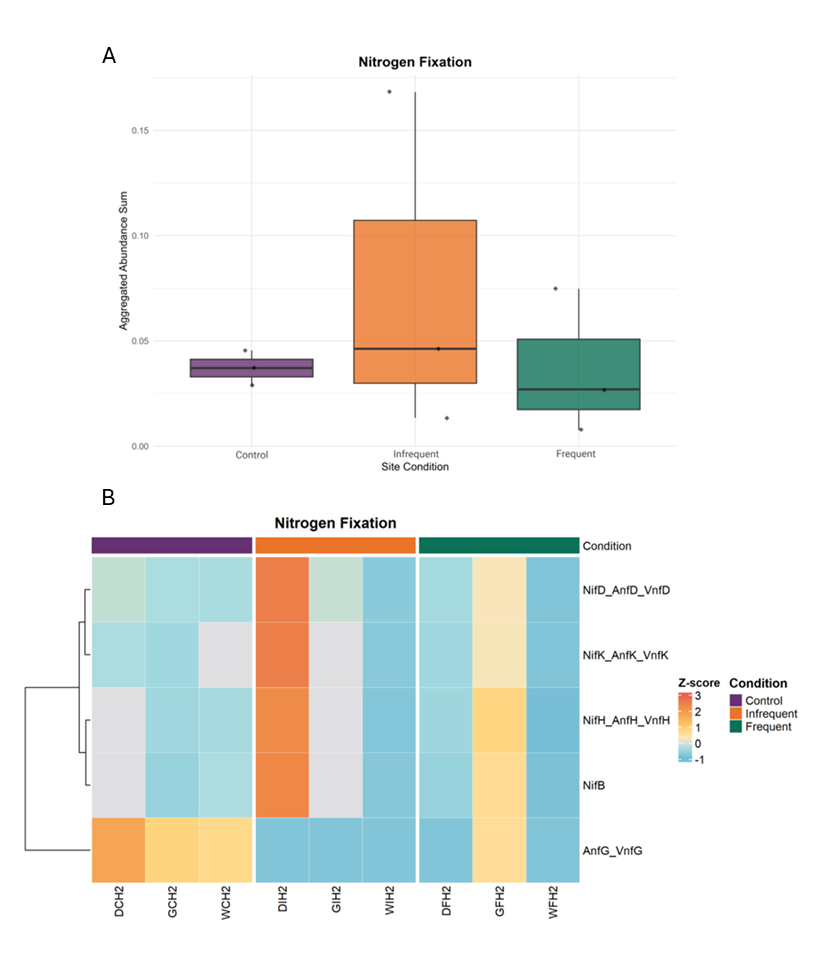

**Figure S2. A)** Normalized heatmap of nitrogen fixation pathway and **B)** box-and-whisker plot of all genes in nitrogen fixation pathway from each site in soil depth 15-30 cm. Data were normalized by column (z-score) and clustered using complete linkage, according to similar distribution patterns.

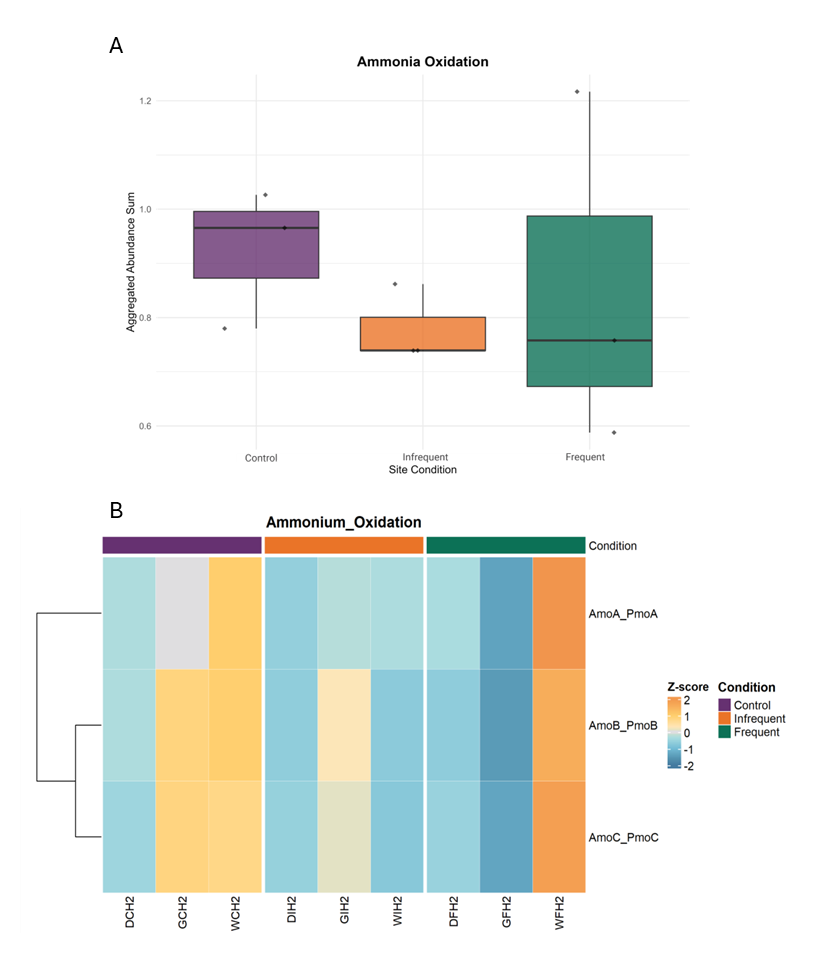

**Figure S3. A)** Normalized heatmap of ammonium oxidation pathway and **B)** box-and-whisker plot of all genes in ammonium oxidation pathway from each site in soil depth 15-30 cm. Data were normalized by column (z-score) and clustered using complete linkage, according to similar distribution patterns.

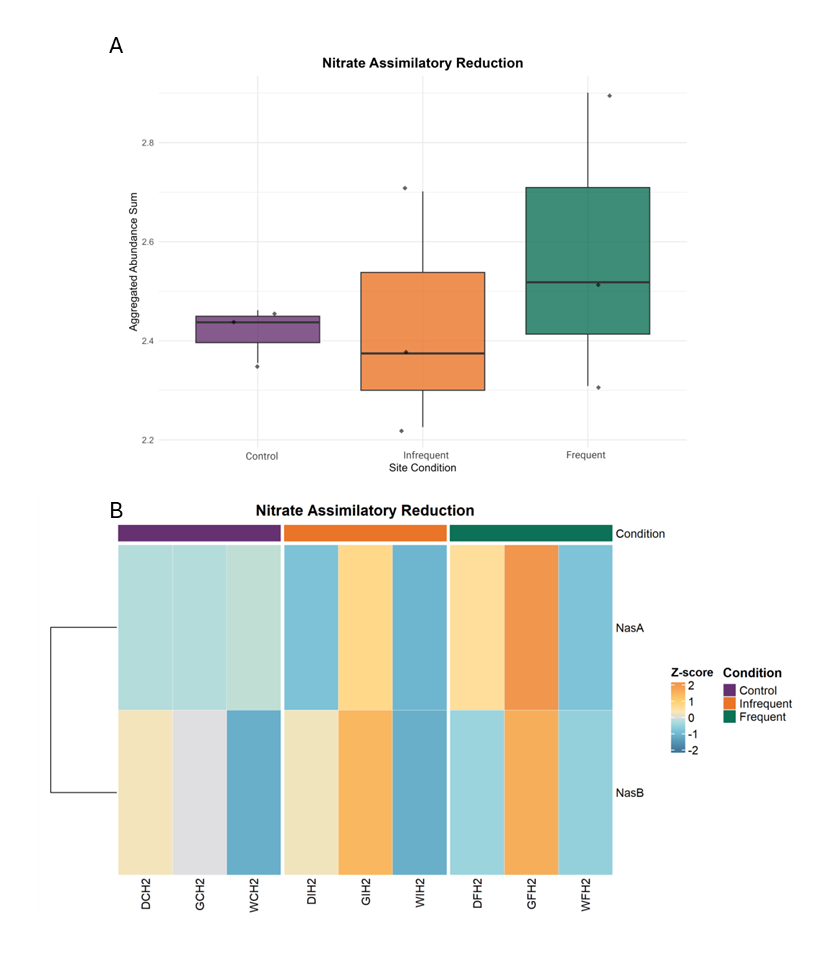

**Figure S4. A)** Normalized heatmap of nitrite assimilatory pathway and **B)** box-and-whisker plot of all genes in nitrite assimilatory pathway from each site in soil depth 15-30 cm. Data were normalized by column (z-score) and clustered using complete linkage, according to similar distribution patterns.

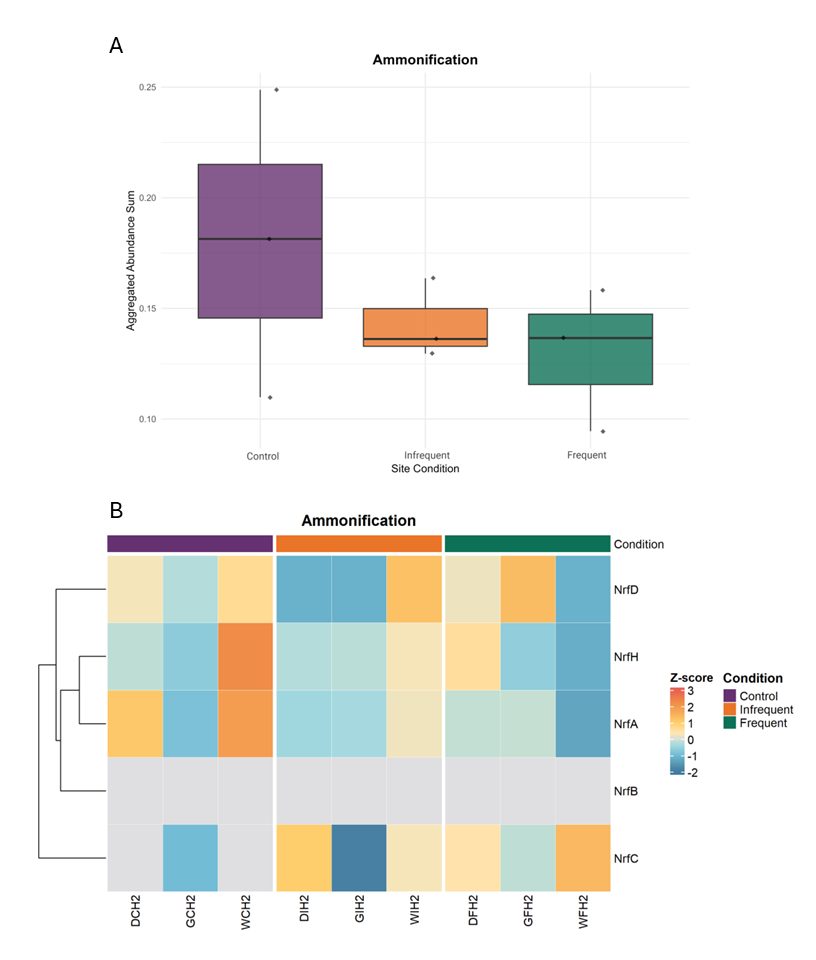

**Figure S5. A)** Normalized heatmap of ammonification pathway and **B)** box-and-whisker plot of all genes in ammonification pathway from each site in soil depth 15-30 cm. Data were normalized by column (z-score) and clustered using complete linkage, according to similar distribution patterns.

B

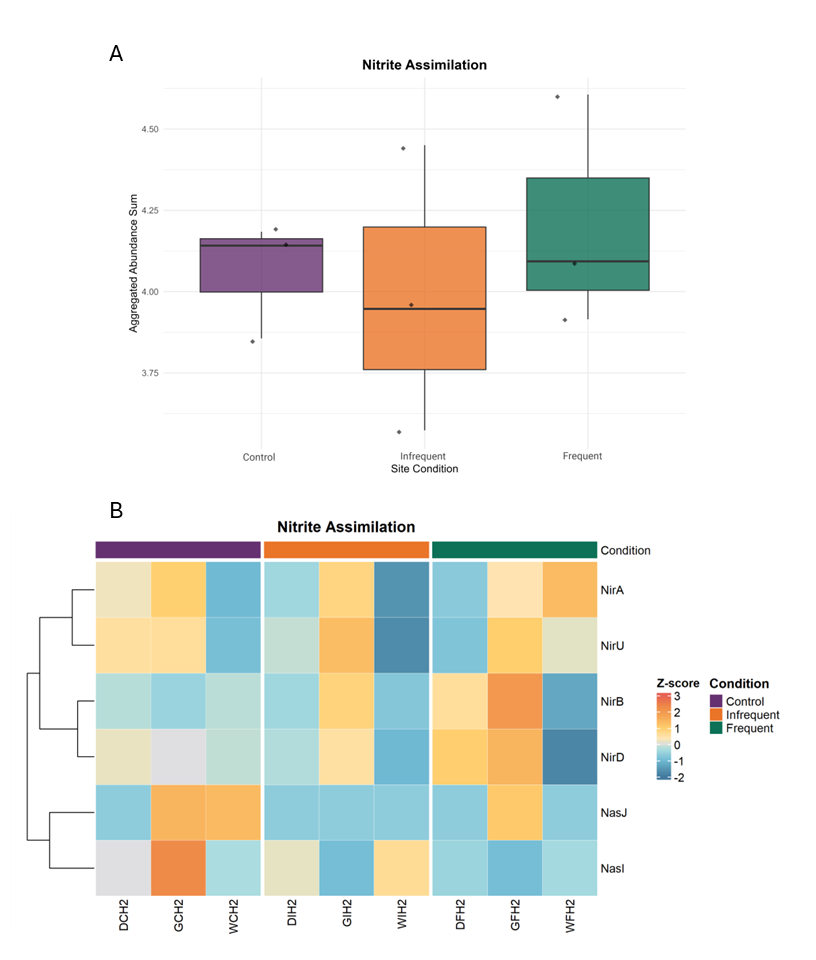

**Figure S6.** **A)** Normalized heatmap of nitrate assimilation pathway and **B)** box-and-whisker plot of all genes in nitrate assimilation pathway from each site in soil depth 15-30 cm. Data were normalized by column (z-score) and clustered using complete linkage, according to similar distribution patterns.

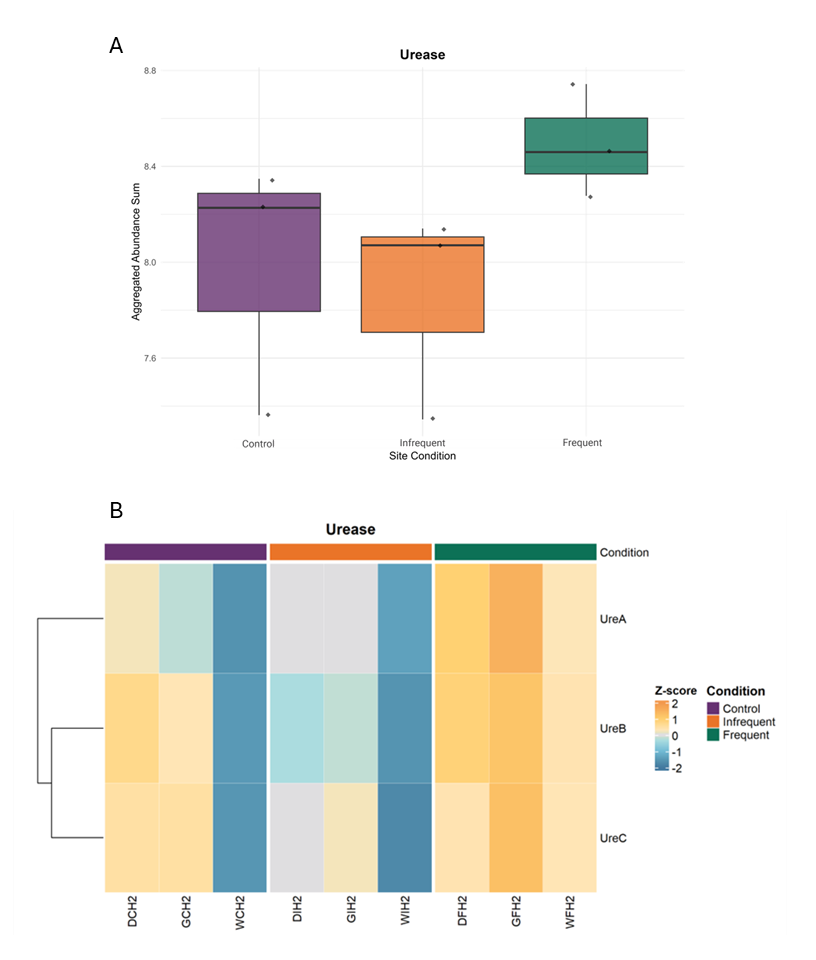

**Figure S7.** **A)** Normalized heatmap of urease pathway and **B)** box-and-whisker plot of all genes in urease pathway from each site in soil depth 15-30 cm. Data were normalized by column (z-score) and clustered using complete linkage, according to similar distribution patterns.

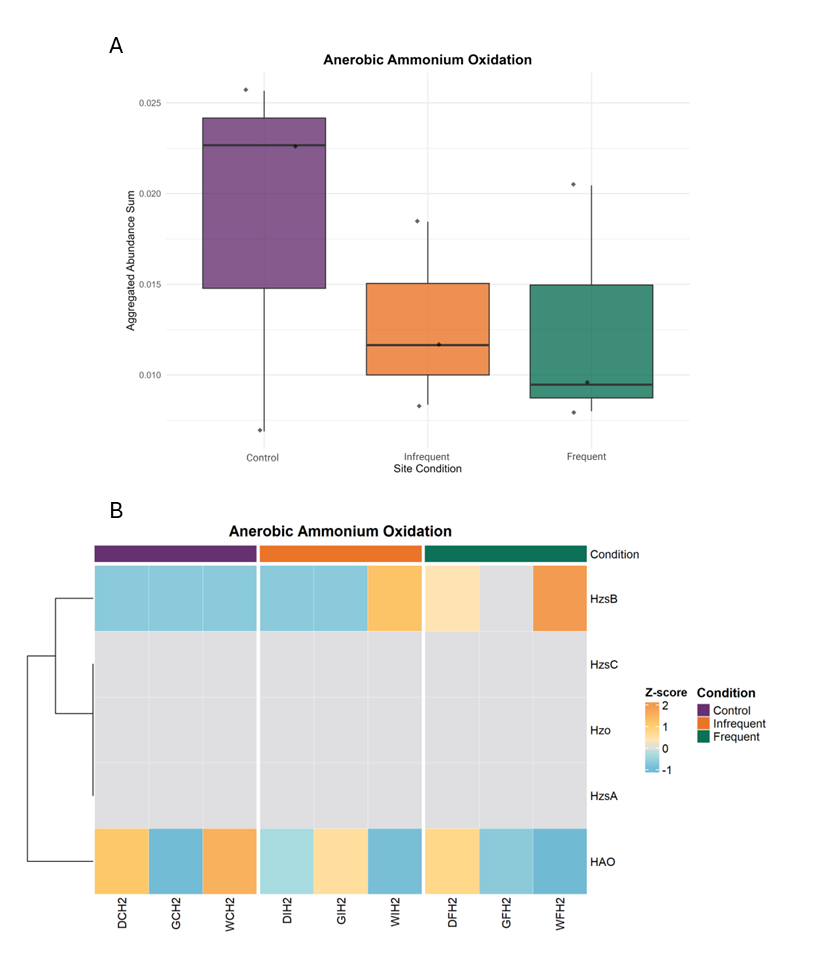

**Figure S8.** **A)** Normalized heatmap of anerobic ammonium oxidation pathway and **B)** box-and-whisker plot of all genes in anerobic ammonium oxidation pathway from each site in soil depth 15-30 cm. Data were normalized by column (z-score) and clustered using complete linkage, according to similar distribution patterns.
